# Predation by migratory birds shapes seasonally fluctuating selection on a conspicuous signal

**DOI:** 10.64898/2026.08.24.746656

**Authors:** C. Patterson, G.F. Grether, F.G. Soley, J. Clavel, M.E. Bonillas Monge, L. Mendoza Cuenca, R. Palin, A. Pérez Madrigal, E.A. Saban-Sequén, A. Tonkinson, J.P. Drury

**Affiliations:** Department of Biosciences, Durham University, Stockton Road, Durham, United Kingdom DH1 3LE; School of Life Sciences, University of Nottingham, United Kingdom, NG7 2RD; Department of Ecology & Evolutionary Biology, University of California, Los Angeles, USA; Escuela de Biología, Universidad de Costa Rica, San José, Costa Rica; Université Claude Bernard Lyon 1, LEHNA UMR 5023, CNRS, ENTPE, F-69622, Villeurbanne, France; Facultad de Biología, Universidad Michoacana de San Nicolás de Hidalgo, Morelia, México; Universidad Estatal a la Distancia, San José, Costa Rica; Facultad de Agronomía, Universidad de San Carlos de Guatemala, Guatemala

**Keywords:** seasonal polyphenism, eco-evolutionary dynamics, species interactions, Odonata, phenotypic plasticity

## Abstract

Sexually selected traits often impose fitness costs on their bearers. Yet, the relative costs and benefits of conspicuous traits can vary through space and time, driving variation in selection acting on those traits. For insects, an important but overlooked source of such variation is seasonal shifts in the local abundance of migratory insectivorous birds. Smoky rubyspot damselflies (*Hetaerina titia*) exhibit a marked seasonal polyphenism in wing pigmentation throughout much of North America, with individuals emerging in the summer exhibiting conspicuous dark wings. Here, we test the hypothesis that this variation is an adaptive response to seasonal and geographical variation in predation risk. First, we find evidence for strong constraints acting on wing phenotypes outside of the summer season, consistent with a seasonal shift in the relative costs and benefits of pigmentation. Second, using a continentally distributed predation experiment, we find that predation risk covaries with spatiotemporal variation in wing pigmentation and is linked to shifts in the local abundance of migratory birds. Overall, our analyses establish an eco-evolutionary link between tropical and temperate regions, underscoring the importance of considering both the evolutionary and ecological consequences of spatiotemporal variation in biotic interactions as species assemblages shift in response to global change.

## Introduction

Conspicuous sexually selected traits often incur a cost in the form of increased risk of being detected by predators or parasites (1, 2). Yet, these costs are not always constant across a species’ range or through time. Across populations, spatial variation in the costs and benefits of possessing conspicuous traits could generate population differentiation via local adaptation (3, 4). Within populations, one way of adapting to shifts in the relative costs and benefits is through polyphenisms, a form of developmental plasticity in which an individual develops one of two or more alternative phenotypes (5). In cases where the relative costs and benefits are temporally variable, populations can exhibit seasonal changes in phenotypic composition (6, 7). Seasonal polyphenisms are hypothesized to evolve when different phenotypes perform best in different seasons due to predictable changes in the environment (7, 8). Caterpillars of the moth species *Nemoria arizonaria*, for instance, develop into catkin mimics in the spring and twig mimics later in the summer, thereby tracking seasonally changing backgrounds such that predation risk is diminished (9).

For insects, migration of insectivorous birds drives predictable seasonal shifts in the local abundance of predators. Each year across the North American continent, bird migration generates massive seasonal fluxes in the abundance of billions of insectivores (10). These dynamics likely generate cyclical patterns of selection on the visual signals of prey. Nevertheless, while shifts in the abundance of migratory insectivores can influence insect abundance (11), the impact of bird migration on insects has received surprisingly little attention relative to the impacts of insects (e.g., phenology) on migratory birds.

Smoky rubyspot damselflies (*Hetaerina titia*) vary both geographically and seasonally in the extent of melanized wing pigmentation (Fig. 1). The seasonal shifts in wing melanization evidently are cued by photoperiod, or an environmental cue closely linked to photoperiod (12). Although melanin deposition occurs during the adult maturation period (∼2 weeks), the area of the wings where melanin is deposited is fixed at adult emergence (12)—in other words, the shift in population means results from a shift in the phenotypic composition of newly emerging individuals. Population differences in both the extent and seasonality of wing melanization are consistent from one year to the next and presumed to be genetic (13, 14). As with other seasonally polyphenic taxa, variation among populations across the species range is thought to represent evolution of the mean and switch point of the developmental reaction norm, not plastic responses to local conditions (6, 7). Populations of *H. titia* on rivers that drain into the Atlantic Ocean (including the Caribbean Sea and Gulf of Mexico) exhibit a pronounced seasonal polyphenism: highly melanized individuals emerge during the temperate region summer (or wet season, in tropical regions) (June-August, the ‘peak’ season) and less melanic individuals, which resemble other sympatric congeners, emerge at other times of year (the ‘off-peak’ season)(13). In populations along Pacific drainages and in the furthest northern reaches of the species’ range, however, the polyphenism is much less pronounced or even absent, with individuals exhibiting overall lower levels of wing pigmentation.

**Figure 1.**
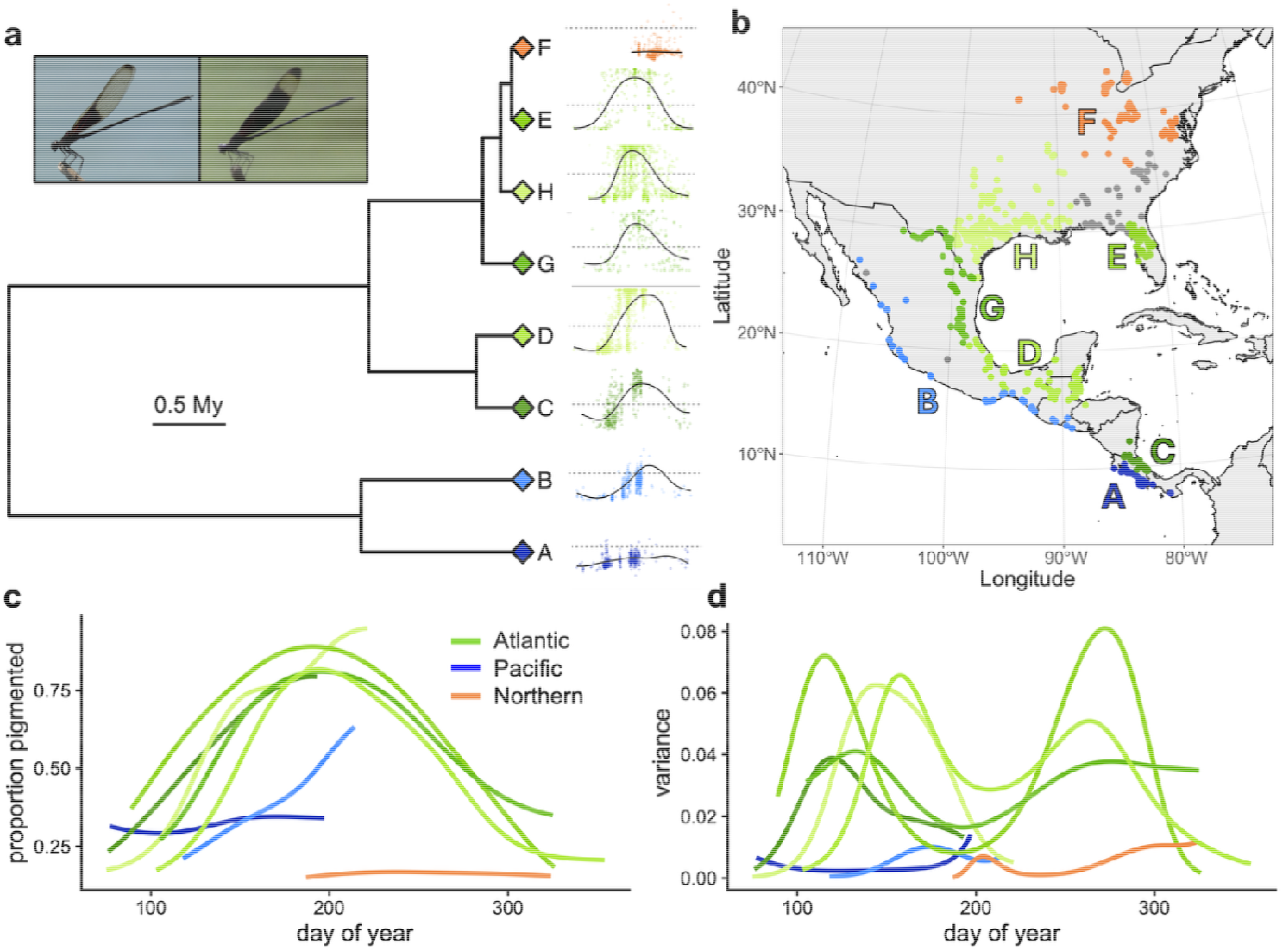
Variation in seasonal polyphenism of smoky rubyspot damselflies (*Hetaerina titia*) across the species phylogeny. **a.**The phylogeny of *H. titia* population clusters with measurements of wing melanization plotted by date of photograph (the dashed line represents 50% of hindwing surface with pigment, the solid line represents model predictions, and the red tick-mark represents the midpoint of the year). The inset photos illustrate variation in the extent of wing pigmentation (photos by G Grether). **b**. Geographic distribution of phenotypic measurements colored and labelled by the assigned tree tip. Grey points mark phenotypic measurements that could not be assigned to the phylogeny due to a lack of genomic data. Model predictions of **c**. mean wing pigmentation in Atlantic, Pacific, and Northern regions and **d**. variance in wing pigmentation for each population cluster, limited to timespans where sampling permitted reliable inference of variance (see Methods).

Wing coloration in rubyspot damselflies is an intrasexually selected trait (15–17) that also strongly influences behavioral interference with sympatric congeners (13–15, 18). In addition to affecting intraspecific interactions, dark wing pigment in *H. titia* males reduces the frequency of interspecific fights and, in females, reduces the rate of reproductive interference compared to non-melanic individuals (13, 14, 18, 19). Consequently, seasonally polyphenic populations of *H. titia* experience high rates of interference with congeners outside the peak of seasonal increase in wing melanization (13), and field experiments show that the shift in wing coloration is largely responsible for the decrease in interference (13).

Given the clear benefits to melanin pigmentation in *H. titia*, it stands to reason that melanic individuals are not ubiquitous because wing melanization is costly. Indeed, dark wing pigmentation increases the conspicuousness and, consequently, predation risk of several related damselflies (20, 21).

Here, we test the hypothesis that the geographical and seasonal variation observed in smoky rubyspot wing pigmentation is an evolved response to spatial and temporal variation in the costs (i.e., predation risk) of possessing conspicuous dark wings. First, by combining field data with automated, computer vision measurements of photographs submitted to iNaturalist, we characterized the seasonal polyphenism across the range of *H. titia* by conducting phenotypic and phylogenetic modelling. Secondly, we carried out a continentally distributed predation experiment to evaluate a possible link between seasonal fluctuations in predation risk and spatiotemporal variation in conspicuous wing pigmentation. Finally, at each site where we conducted the predation experiment during the spring bird migration period, we extracted the modelled weekly relative abundance of migratory invertivores from eBird Status & Trends (22) to test whether shifts in migratory bird dynamics influence the relative costs of possessing dark pigmentation.

## Results

### Constraints on wing pigmentation shift seasonally

Using whole-genome sequences from 33 individuals, we reconstructed a phylogeny of smoky rubyspot population clusters for which we could characterize the seasonal polyphenism using a phenotypic dataset from >5800 males (Fig. 1a,b, Supplementary Table 1, Supplementary Figures 2-4). As determined previously (23), we found that the seasonal polyphenism is distinct in three different regions, with the most pronounced shift in wing coloration occurring in population clusters in Atlantic drainages along the Caribbean and Gulf of Mexico (Fig. 1c, Supplementary Tables 3-5). Similarly, the Pacific and Northern regions exhibit low variance in wing pigmentation throughout the year, but the variance in the Atlantic tends to be lowest in the off-peak season (i.e., when *H. titia* wings exhibit the lowest amount of melanin pigmentation) relative to the peak season (i.e., when wings exhibit the highest levels of pigmentation), with the highest variance observed during transitions between the two (Fig. 1d, Supplementary Table 5).

Modelling the evolution of peak and off-peak phenotypes using phylogenetic models, we found that a model of constrained evolution (i.e., Ornstein-Uhlenbeck) was the best-fitting model for both peak and off-peak phenotypes (Supplementary Figures 4-5, Supplementary Table 6). However, the stationary variance—a measurement of the degree of constraint estimated from the model—was > 20x smaller in the off-peak season than in the peak season, consistent with a stronger evolutionary constraint acting on off-peak phenotypes (Supplementary Table 6).

### Predation risk drives seasonal shifts in wing pigmentation

To test the key assumption of the predation-cost hypothesis—that is, that highly melanized wings are more conspicuous than wings with low levels of melanization (24)—we extracted measurements of conspicuousness from perched male and female individuals and found that individuals with more pigmentation were indeed more conspicuous (Supplementary Figure 7; males: conspicuousness∼proportion of wing pigmented *t* = 11.948, *p* < 0.001, Adj. R^2^ = 0.73; males: conspicuousness∼wing illuminance *t* = -8.426, *p* < 0.001, Adj. R^2^ = 0.57; females: conspicuousness∼wing illuminance *t* = -7.548, *p* < 0.001, Adj. R^2^ = 0.75).

We tested the prediction of the predation-cost hypothesis, that predation risk would be lowest at locations and times of the year where wing melanization is highest, using a continentally distributed predation experiment with >1500 plasticine damselfly replicas (Fig. 2a-c). Overall, we found striking support for the predation-cost hypothesis: predation risk was highest in the Pacific drainages, which exhibit relatively low levels of wing pigmentation year-round, with no differences between predation risk in spring and summer in this region (Fig. 2e, Supplementary Tables 8-9). In the Atlantic region, predation risk was higher in the spring, when individuals exhibit low levels of wing pigmentation, than in the summer, when wing melanization is at its maximum level. The exception was in the northern drainages, where predation risk was very low despite low levels of melanization. The best-fitting model, which included the region X season interaction, was within 2 AIC units of a model without the interaction (Supplementary Tables 8-9), but the differences between seasons and regions were consistent with the predation-cost hypothesis in both models.

**Figure 2.**
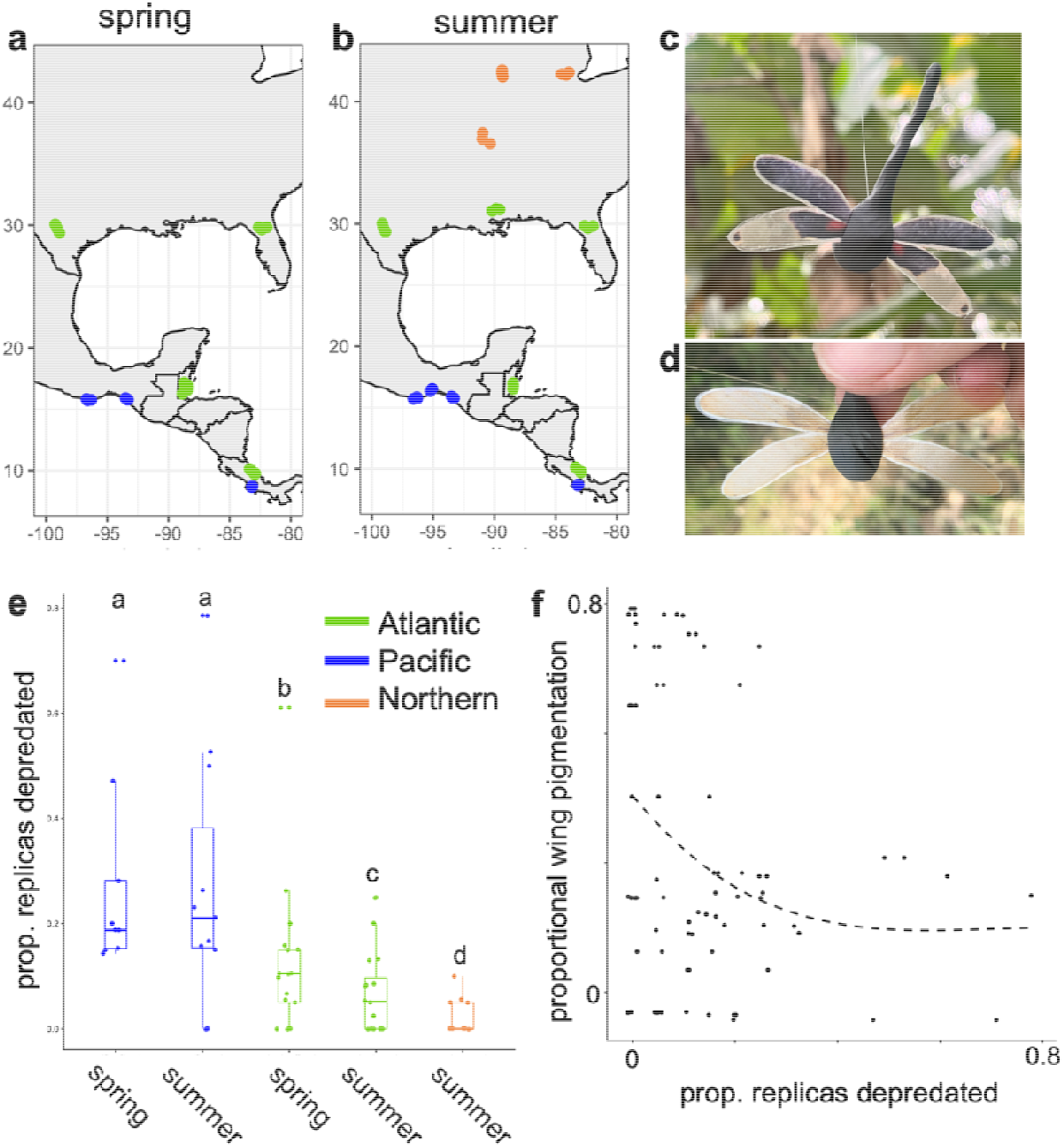
A continentally distributed predation experiment supports the predation-cost hypothesis. **a-b.**Sites where predation experiments were deployed during the spring (*n* = 34) and summer (*n* = 42). **c**. An example of a plasticine damselfly replica *in situ* and **d**. an example of a model with a beak imprint demonstrating a predation attempt (see also Supplementary Figure 8). **e**. Predation risk was highest in Pacific drainages throughout the year, and then highest during the spring in Atlantic drainages, with Northern drainages experiencing overall low predation risk (the letters depict statistical differences between groups; note, experiments were not conducted during the spring for northern populations because adult damselflies emerge later in the year). **f**. The proportion of damselfly replicas attacked was negatively correlated with the proportion of hindwing with pigmentation expected during the deployment of predation experiments (the dotted line is a loess curve fit to the data). In both **e** and **f**, points are jittered horizontally for visualization.

When directly comparing the predation rate to the predicted local mean wing phenotype (Fig 1a) at the site and time of plasticine model deployment, we found that the mean wing pigmentation decreased as predation risk increased (Fig. 2f; Spearman’s rank correlation: mean proportion pigmented vs. proportion attacked ρ = -0.23, *p* = 0.048).

### Migratory fluxes in insectivorous birds influence shifts in predation risks

There was a clear relationship between predation risk for damselfly replicas and bird migration. Accounting for the overall higher predation risk in Pacific drainages, predation risk was highest at the time of peak migration, declining with time from the week of peak relative abundance of migratory insectivores (Fig. 3a, Supplementary Table 10). Similarly, as the magnitude of the shift in relative abundance of migratory birds from the time of deployment to the non-migratory period increased, predation risk increased in Atlantic drainages (Fig. 3b, Supplementary Table 11). In the Pacific, there was a negative relationship between the magnitude of the shift in relative abundance and predation risk (Fig. 3b), but this relationship was not strongly supported; a model without an interaction term (and a positive relationship) was within 2 AIC units of the interaction model (Supplementary Table 11). These patterns were consistent when limiting analyses to clear bird attacks and likely bird attacks (Supplementary Table 12).

**Figure 3.**
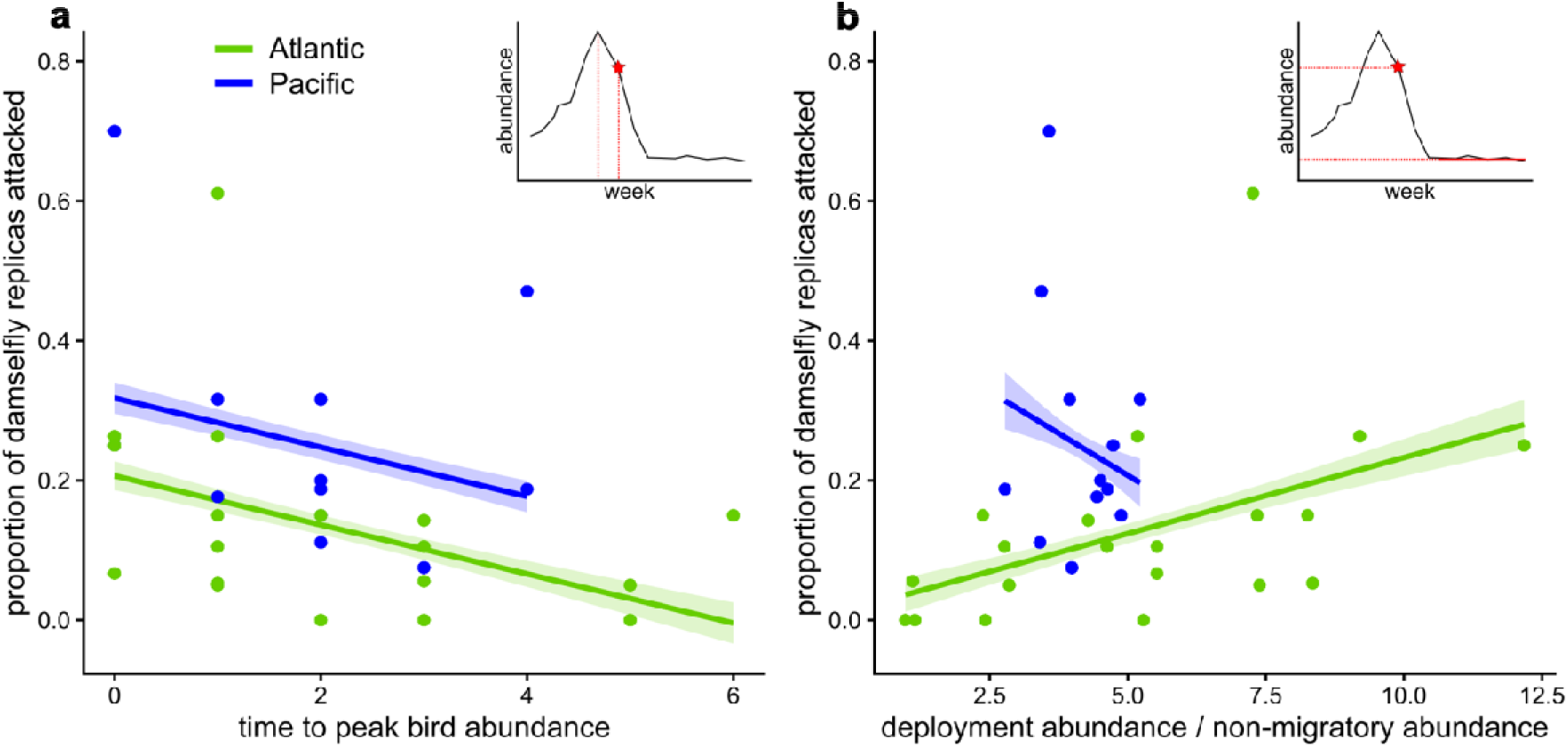
Predation risk covaries with migration of insectivorous passerines. **a.**Predation risk declines as the time (in weeks) between the deployment of the damselfly replicas and the migration peak (modelled using eBird Status & Trends data) increases. **b**. In the Atlantic, predation risk increases with the magnitude of the migratory flux, measured as the relative bird abundance at the time of deployment divided by the relative bird abundance in the non-migratory period (late May to July), but in the Pacific, this relationship is reversed. The inset plots depict a relative abundance-through-time curve, with stars representing the deployment of the damselfly replicas and lines representing the values being compared in each index.

## Discussion

We found that variation in predation risk predicted both temporal and geographic variation in wing pigmentation in smoky rubyspot damselflies throughout much of their range, and that this predation risk covaried with the shifting abundances of migratory insectivorous birds. Previous research has demonstrated a relationship between predation risk and conspicuous signaling traits (3, 20, 21, 24) and a relationship between predation and polyphenism, including seasonal polyphenism (25–27). However, to our knowledge, ours is the first study to link the evolution of a seasonal polyphenism to the predictable annual shifts in the abundance of insectivorous birds that result from bird migration. Migratory flyways in North America have existed for millions of years (28), and annual fluxes in the abundance of migratory birds likely have important ecological consequences for the local communities along their migratory paths (29). Our work demonstrates that migratory dynamics also shape dynamics in selection pressures acting on prey species, imposing a trade-off in sexually selected signaling.

Nevertheless, spatiotemporal variation in predation risk did not universally covary with wing pigmentation—dark wing pigmentation is the least pronounced in the northernmost populations, despite a much lower estimated predation risk. The northern part of the range of *H. titia* (Fig. 1b) must have been colonized from the south sometime after the end of the Pleistocene glaciation, 11,700 years ago (30), meaning the reduction in wing melanization in the north is clearly a recent evolutionary event. Whether this reduction results from costs other than predation risk (e.g., physiological costs associated with pigment production, behavioral interference with North American *Calopteryx* damselflies) or differences in the benefits of wing pigmentation is currently unknown. Another possibility is that predation attempts have higher success rates in the northernmost populations, owing to damselflies’ reduced ability to warm their flight muscles due to lower air temperatures in the early part of the day.

We also found that the times of the year when predation risk was highest corresponded to the highest levels of constraint acting on wing pigmentation estimated from phylogenetic models of phenotypic evolution. This is consistent with selection for low levels of wing melanization in the spring being stronger than selection for high wing melanization in the summer. Relatively weak selection in the summer may indicate that the benefits of wing melanization diminish as proportional wing melanization increases (19). In aggregate, our results support the hypothesis that a trade-off between predation and competition drive evolutionary dynamics of a reaction norm shaping the mean, as well as the variance, of wing pigmentation.

The link between predation risk and the evolution of a seasonal polyphenism provides further evidence of the role that trophic interactions can play in structuring evolutionary dynamics in signaling traits. Moreover, our results reveal how transient trophic interactions between temperate and tropical species can shape evolutionary trajectories in tropical species. As species assemblages shift in response to rapid changes in climate and land-use, the presence and magnitude of biotic interactions are likely to change, as well (31). Indeed, migratory flyways are likely to experience restructuring as a result of ongoing human-driven global change (32). As with other global-change driven restructuring of trophic communities (e.g., plant-pollinator interactions (33)), shifting assemblages may lead to changes in the selective trade-offs that shape intraspecific interactions. Future research will be critical to characterize not only the ecological, but also the evolutionary consequences of changing biotic interactions.

## Materials and methods

### Measurements of wing pigmentation

Both female and male *H. titia* exhibit variation in the extent of wing pigmentation. Male *H. titia* have red basal wing spots, similar to other rubyspot damselflies (*Hetaerina* spp.), but the red basal colour on the hindwings can be fully or partially masked in highly melanized individuals. Male *H. titia* defend territories along riverbanks while females spend a high proportion of time foraging at higher levels. Consequently, male *H. titia* are substantially easier to photograph (>80% of iNaturalist photos of *H. titia* are male), and, as such, we limited our analysis to male phenotypes.

We measured the relative size of pigment patches on male wings from photographs obtained from two sources. Firstly, a photographic dataset of *H. titia* was collected between 2004 and 2024 during scientific fieldwork (*n* = 3956; see methods in ref (23)). Additionally, to increase the spatial and temporal scale of our dataset, we also obtained images from the citizen science platform iNaturalist (https://www.inaturalist.org). We downloaded images of *H. titia* submitted to iNaturalist prior to 2024. We manually curated these photos to only include perched adult male *H. titia* with the entire wing visible within the photo (*n* = 1936).

For many iNaturalist photos, we manually measured the proportion of hindwing with pigment for each photo in ImageJ (34) using previously published methods (23). Additionally, we developed an automated image detection and segmentation pipeline for automated extraction of wing pigmentation estimates from iNaturalist photos. The automated image detection pipeline uses Detectron2 (35) to segment out each wing and remove the background from each image. The segmented wing was then analyzed using OpenCV (36) to detect the proportion of pigmented pixels using manually defined threshold ranges in HSV and RGB colour spaces. Each automatically segmented wing and extracted pigment patch was manually inspected for errors, and measurements from photos with errors were removed. The full automated image detection pipeline is available at https://github.com/ChristophePatterson/wing_damselfly.

To increase the temporal resolution of the dataset, we first identified date ranges at the beginning and end of the year with few observations (< 20 per two-week window). We then targeted iNaturalist observations from these time windows that were submitted between 1 Jan 2024 and 11 Feb 2026 for measurement. In cases where we found unreliable segmentation or patch extraction based on the automatic pipeline, we made manual measurements.

We then standardized all measurements to ensure analyses are robust to differences in data source or measurement strategy (Supplementary Methods, Supplementary Figure 1).

### Phylogenetic reconstruction

We reconstructed the phylogeny of *H. titia* using SNP data from whole-genome sequencing (Supplementary Table 1). The raw sequences were mapped to the HetTit1.0.p draft genome of *H. titia* (37) using bwa 0.7.17 (38). All samples met our minimum quality threshold, having a mean coverage above 5X and a mean mapping QC above 10. Duplicate mapped reads were removed using the MarkDuplicates function in picard (39). SNP calling was conducted using bcftools (40), using the mpileup and call functions. In mpileup, we set a minimum read mapping threshold of 20 and a minimum base quality of 13. We removed indels, set SNP calls with a depth less than 3 to missing, removed SNPs with an average depth below 3 and above 1.5x the mean coverage (≤ 27.42x), removed all non-biallelic SNPs, and then filtered to only include SNPs with 50% coverage. Following this, to avoid non-independence owing to linkage disequilibrium, the -thin function from vcftools (41) was used to create a vcf file containing only the first SNP within 10,000 bp windows.

To place our samples in the geographical context needed to identify regions for which we could estimate the seasonal shift in wing pigmentation (Supplementary Methods), we assigned samples to the 5^th^ level of the hydroBASIN dataset (42) using the function *st_intersection* within the R package *sf* (43). The hydroBASIN dataset is a global model of river basins consisting of 12 levels of hierarchical nested sub-divisions allowing river catchments to be subdivided into consistently sized basins. We identified the highest-coverage samples from each level 5 hydroBASIN with samples, removed SNPs that were either no longer polymorphic between the selected samples or not genotyped across all individuals, yielding 42,313 polymorphic SNPs across 33 samples.

To reconstruct the phylogeny, we ran the Bayesian coalescent analysis SNAPP(44) implemented within the programme BEAST v2.7.5(45). A SNAPP configuration file was created using a custom R script and the ruby script from https://github.com/mmatschiner/snapp_prep. We set SNAPP to sample every 500 iterations. To time-calibrate the tree, we used a previous estimate of the crown ages of *H. titia* to select priors (mean = 3.7 million years ago (Mya), standard deviation = 0.1 million years)(46).

Convergence was assessed using Tracer, and the maximum clade credibility tree, with the first 20% of the posterior sampled removed as burn-in, was calculated using the standard settings of the TreeAnnotator program distributed with BEAST.

*H. titia* is currently recognized as a single highly variable species, but population genomic analyses suggest that the Pacific and Atlantic populations diverged ∼3.7 million years ago and that there is reproductive isolation at a site of secondary contact (46). The taxonomic classification of *H. titia* requires further investigation but does not influence our inferences about the evolution of the polyphenism.

### Phenotype modelling

We modelled the evolution of the seasonal polyphenism at the highest resolution that the phylogenetic reconstruction allowed (see Supplementary Methods). As such, we used the highest resolution phylogeny of *H. titia* for which we could estimate the level of melanization throughout the year for each tip (see Supplementary Methods).

Using the R-package^45^ brms (47), we modelled the annual dynamics in the proportion of wing pigmentation for each of the eight population clusters included in our phylogenetic reconstruction as a smoothed function (*k* = 8) of the day of the year of each observation (with separate smooth terms estimated for each cluster to estimate different curves for each cluster). Given the large number of observations with 100% hindwing pigmentation, we used a zero-one-inflated beta family. We fitted both the precision (*φ*) and zero-one inflation terms as smoothed functions (*k* = 8) of the day of year, again with separate smooth terms estimated for each cluster. We also included a random effect for site, rounding each observation’s latitude and longitude to the nearest tenth of a degree (∼11km), to address spatial variation in sampling. For all smooth terms, we used a cyclic cubic spline to model the annual cycle (with knots at 1 and 365). The model fit was conducted via MCMC sampling of four chains, each run for 3,000 iterations with a warm-up period of 1,500 iterations. Convergence was assessed using Rhat diagnostics (all Rhat values =1-1.01). This model outperformed simpler models without cluster-specific smooth terms (Supplementary Table 4).

Although the above beta regression explicitly models variance, we found that estimation of the variance in the above model was biologically unrealistic due to poor estimation of *φ* in clusters and at times of the year with poor sampling. Consequently, we focused our analysis of variance over the year to a model that was fitted to a dataset restricted to the time window starting in the first two-week period with at least 10 observations and ending in the last such period (*n* = 5650; Supplementary Table 5). Otherwise, this model was identical to the model with all data. Variance was calculated under the beta family as μ * (1 - μ) / (1 + *φ*).

To estimate the tempo and mode of wing phenotype evolution across the population-level phylogeny, we used the R-package rphylopars (48) to fit Brownian motion (a drift-like model where traits diffuse through trait space over time) and Ornstein-Uhlenbeck (OU; a model including a constraint parameter) models to our dataset, incorporating individual measurements of wing pigmentation across the eight population clusters represented in our population-level phylogeny, separately for peak (i.e., within two-weeks of the maximum predicted mean from the main beta regression above, ranging from 26 June to 1 September) and off-peak (between 5 November and 25 April) measurements (see Supplementary Methods, Supplementary Table 6, Supplementary Figure 4). We transformed proportional wing pigmentation (*p*) using a logit transform:

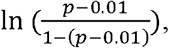

to conform to model assumptions. We verified that patterns were similar across different definitions of ‘peak’ (28-day and 14-day windows) and conducted simulation and model adequacy tests to explore the robustness of inferences under our modelling framework (see Supplementary Methods, Supplementary Figure 5).

### Conspicuousness of melanized wings

We photographed a sample of males (*n* = 53) and females (*n* = 20), spanning a range of wing melanization phenotypes (Supplementary Table 7). Each individual was photographed in two ways: (1) against a background of riverbank vegetation with a Nikon D7000 DSLR (18-55mm lens), with a colour standard in each photo (ColourChecker Passport, Calibrite) and (2) with a standardized (white) background with each wing separated. RAW format photo files were calibrated using the generate multispectral image function in the MICA image analysis toolbox plugin (49) with NIH ImageJ(34), setting the standard reflectance of the visible patches of the grayscale colour standard, which were determined using reflectance spectroscopy (Ocean Optics Flam-S-UV-VIS spectrometer equipped with a pulsed xenon light source (Ocean Optics PX-2)). The hindwing surface was then outlined in each perched photo, and the mean luminance was calculated using batch multispectral image analysis. We then calculated conspicuousness as the just noticeable differences (JNDs) between the outlined wing and a similarly sized background selection in each perching photo (5). To test for a relationship between wing pigmentation and conspicuousness, we fitted linear models comparing conspicuousness to the overall luminance of the hindwing surface (from the standard photos) for both sexes and also the proportion of pigment in the male wings (measured from the perching photos as in reference (23)).

### Predation experiment

At 42 streams distributed from Costa Rica to the northern United States where *H. titia* has been observed (Fig. 1a,b), we hung twenty plasticine damselfly replicas to vegetation overhanging the stream. The body of the damselfly replicas was made of wire covered with black plasticine and the wings were made from polyester film imprinted with images of *H. titia* wings. We grouped these replicas into four ‘stations’, where each station comprised five replicas. Each such station was separated from other stations by a minimum of 20 meters. Within each station, we ensured that replicas were in the same general area (i.e., w/in 4m^2^), a minimum of 30cm separating each treatment. The replicas were placed between chest-height (∼1.3m) and arms-length (∼ 2.5m) height such that replicas were visible from >3m away (i.e., where they were not hidden by the leaves of the tree to which they were attached), within daylight hours.

During each deployment of damselfly replicas, we checked them three times: after 24h, 48h, and 96h (4 days) ± 2 hours from the time they were first placed in the field (following (50)). Damselfly replicas that had been attacked were removed and photographed. If a replica could not be found, we recorded it as ‘missing’. Neither attacked nor missing replicas were replaced. Where possible, we deployed damselfly replicas multiple times at each site (mean number of deployments = 1.8, range = 1-5). In total, we conducted 76 deployments, comprising 1515 damselfly replicas.

We classified the source of each attack according to the imprint on the plasticine replicas, using a combination of field notes, drawings and posterior re-examination of photos by JPD and GFG. We classified obvious beak marks as ‘clear bird’ attacks (58.4% of attacks), cases where sharp clean lines seemed most consistent with beaks but not unambiguously so as ‘likely bird’ attacks (11.4% of attacks), and others as ‘other’ (30.2% of attacks, most of which were made by lizards). Examples of each of these attacks are presented in Supplementary Figure 8.

### Bird migration data

We obtained data on the timing of avian migration at each site where predation experiments were deployed using eBird Status and Trends (22). In the R package ebirdst (51), we subset data for North American migratory insectivorous passerines (*n* = 156 spp.), using the 2022 data version. For each site where we deployed the predation experiment during the spring (i.e., March and April), we extracted the mean modelled relative abundance for each species from within a 10km radius of our study site for each week throughout the year and summed this value across all species. Using these data, we identified the week with the maximum relative abundance of migratory invertivores between the last week of February to mid-May and determined the number of weeks separating our deployment from the week of peak insectivore relative abundance in this period. Similarly, as a measure of the relative magnitude of the shift in abundance in migratory species, we calculated the ratio of the relative abundance at the time of deployment to the relative abundance during the ‘trough’ in migrant abundance (between late May and the last week in July).

### Predation risk analyses

To estimate predation risk, we fitted multi-state Markov models in the R-package msm (52). To test hypotheses about the factors that influence predation risk, we compared the fit of null models—that is, models where the transition rate from ‘not predated’ to ‘predated’ was constant—to models where transition rates vary as a function of covariates using AIC for model comparisons. Damselfly replicas that were missing were treated as censored data.

To directly relate the predation risk to the wing phenotypes of *H. titia* present at the time of deployment, we used the model of annual variation in wing pigmentation (see *Phenotypic modelling*, above) to predict the mean and variance in the proportion of wing pigmentation at the time of deployment (i.e., by calculating the mean of the predicted mean and variance for the week of the deployment). We then tested for a correlation between predicted mean and variance values and the proportion of damselfly replicas attacked in each deployment using Spearman’s rank correlation tests.

## Acknowledgements and funding sources

This research was funded by NSF DEB-NERC-2040883 to JPD and GFG. Specimen collection was conducted under permit SGPA/ DGVS/04421/21 issued by the Mexican Secretaría de Medio Ambiente y Recursos Naturales (SEMARNAT) to LMC; permit 08112112 issued by the Florida Department of Environmental Protection to JPD; license FD/WL/7/21(04) issued by the Forest Department in Belize to JPD & GFG; In Costa Rica, research was performed under permits SINAC-ACOSA-DASP-PI-INV-046-2021, M-P-SINAC-PNI-SE-013-2022, M-P-SINAC-PNI-SE-004-2023, SINAC-ACOPAC-D-RES-041-2023, and M-P-SINAC-PNI-SE-004-2024 from the Sistema Nacional de Áreas de Conservación (SINAC, MINAE), permit R-018-2025-OT-CONAGEBIO from the Comisión Nacional para la Gestión de la Biodiversidad, and genetic access permission No. 377 issued by the Comisión Institucional de Biodiversidad, Universidad de Costa Rica. We thank M de Jong and J Lin for bioinformatics assistance and M Springer for assistance with permitting in Costa Rica. We thank L Davis and AE Finneran for assistance producing damselfly replicas and K Acuña, K Angulo, S Barboza, R Barajas, V Liseth Calero, J Camargo, J Carballo, J Cerdas, J Obando Chavarría, H Cowling, AE Finneran, R Gil, P Gutiérrez, E Soto Guevara, E Rodríguez, and R Yaxhá for field assistance. For measuring damselfly images and assistance with data curation, we thank YB Abu Ghalyoun, V Agarwa, JH Ahdoot, XA Buschbacher, SC Balotescu, L Boiton-Rodriguez, Y Chen, AE Finneran, AYA Fu, AN Gill, A Grove, M Huang, VS Iyer, MA Lynch, ML Mattis, EWW Meglah, AM Mele, AD Phillips, AA Postajian, J Roshala, AV Scott, E Scholes, EW Sun, S Wolin, and RM Yang. We also thank iNaturalist users for posting their photographs of rubyspots.

## Notes

### Competing Interest Statement

The authors have declared no competing interest.

### Summary of Updates

This manuscript has been revised to clarify a few points, improve figures, and update some supplementary tables.

