## Supplementary Information for "Predation by migratory birds shapes seasonally fluctuating selection on a conspicuous signal"

**Supporting Information for “Predation by migratory birds shapes seasonally fluctuating selection on a conspicuous signal”**

Patterson, C., Grether, G.F., Soley, F.G., Clavel, J., Bonillas Monge, M.E., Mendoza Cuenca, L., Palin, R., Pérez Madrigal, A., Saban, E., Tonkinson, A., & Drury, J.P.

Supplementary Text

Figs. S1-S8

Tables S1-S13

**Supplementary Text**

**Phenotypic data standardisation.** To ensure standardisation between all three of the wing measurement methods, we collected a sample of images of the same individual under both standard and perched conditions (the latter being representative of photos uploaded to iNaturalist). We then used general additive models (GAMs) to model the standardised manually-obtained wing pigmentation measurements as a function of manual measurements of perched photos (*n* = 66) or automated measurements of perched photos (*n* = 115) from the same individual (Supplementary Table 2, Supplementary Fig. 1), using the gam function of the R package mgcv(1), setting the smoothing parameter to k=4.

In our full dataset, to standardise measurements obtained from different data sources, we transformed manual and automated measurements from iNaturalist observations onto the scale of standard measurements using predictions of the GAM for the corresponding measurement methodology.

**Phylogenetic analyses.** To model the seasonal polyphenism at the highest possible spatial resolution, we first collapsed nodes in the population-level phylogeny that were not well resolved (posterior probability <0.9) in the phylogeny (Supplementary Figure 2). We then sought to manually identify monophyletic clusters with sufficient phenotypic data to model the annual trajectory of wing pigmentation, assigning hydrological basins to the nearest sample location and the corresponding tip of the phylogeny (Supplementary Figure 3). One well-supported node (splitting populations on the Pacific coast of Mexico at the Isthmus of Tehuantepec) did not contain enough phenotypic measurements to reliably model the seasonal polyphenism separately in each daughter lineage, so we collapsed this node for subsequent analyses.

The peak level of melanization does not occur at the same time of year for all populations of *H. titia*. The first and last emergence of *H. titia* also varies geographically. As such, we calculated the time of the peak phenotype as the Julian date of the maximum predicted value under our best-fitting model (Fig. 1a, Supplementary Table 3). For two regions (“A” and “F”), the fitted curves were not unimodal and did not have a peak pigmentation value in the summer (Fig. 1a), so we used the mean peak date across other regions as the peak season date. We considered all observations within two weeks of the peak date to be ‘peak’ season observations and calculated the mean and standard error of the logit-transformed wing pigmentation. To verify that the window size did not influence our conclusions, we repeated analyses limiting observations to those within one week of the peak date.

There were fewer observations in the early and late parts of the year, which likely reflects generally lower densities outside of the summer. To calculate an off-peak value for wing pigmentation, therefore, we first calculated a ‘global’ peak season date from all combined observations. Then, we set this peak date (2 August) as the middle of the ‘peak-half’ of the year and went on to calculate an ‘off-peak half’. Setting a two-week buffer period between the halves, this left an off-peak period from day of the year 312 to 115 (Supplementary Figure 4). As above, we calculated the mean and standard error for all observations in this time period.

**Simulation and model adequacy tests.** The distribution of wing pigmentation during the peak season was not normally distributed in several of the clusters (Supplementary Figure 4). To determine whether these distributions influenced our ability to differentiate between BM and OU models, we conducted a simulation analysis. Using the maximum likelihood parameter estimate for $\sigma$^2^ from the BM model fit in Rphylopars, we simulated 2,500 datasets. For each simulation, we then generated a dataset for each tip (of the same size as the sample size for that tip in the empirical dataset) with the simulated value as the mean and the variance, skewness, and kurtosis matching the empirical distribution (using the R package moments(2)). We then fitted both BM and OU models to these BM-generated datasets and conducted model selection as in the empirical analysis. Of the simulated BM-generated datasets, we found BM was selected in the overwhelming majority of the cases (96.64%), demonstrating that the non-normality in our tip distributions was not a viable explanation for OU being the best-fitting model.

To determine whether the best-fitting OU models were compatible with our empirical dataset, we conducted model adequacy tests. We simulated 2,500 datasets under maximum likelihood parameter estimates for the OU models fit to both peak and off-peak data. We then examined how closely the empirical peak and off-peak data corresponded to the coefficient of variation of the absolute value of the contrasts (C_VAR_) and the slope of a linear model fitted to the log-transformed absolute value of the contrasts against node depth (S_HGT_, also commonly referred to as the ‘node height test’)(3). Finally, we compared the disparity through time of simulated datasets to our empirical datasets using the rank-envelope test(4). Differences between simulated and empirical values for either of these indices would suggest that rate heterogeneity was not adequately captured in our model fit. For all indices, our empirical datasets fell within the middle 95% of the distribution of the simulation datasets (Supplementary Figure 6), indicating that the OU model does an adequate job of capturing the variation present in our empirical datasets.

**Table S1.** Sample information for specimens used in reconstructing the population level phylogeny of *Hetaerina titia*.

| **Sample Name** | **Tip Label** | **Latitude** | **Longitude** | **Sex** | **Collection Date** | **NCBI**  **Accession** |
| --- | --- | --- | --- | --- | --- | --- |
| HCARa04 | E | 27.88 | -82.17 | M | 07/08/2019 | SAMN55000492 |
| ZANAa05 | B | 16.45 | -94.34 | M | 13/06/2021 | SAMN55000530 |
| HXRCb03 | B | 15.75 | -96.3 | F | 08/06/2021 | SAMN55000468 |
| CUAJa03 | B | 16.79 | -95.01 | F | 22/06/2021 | SAMN55000462 |
| NR02.10 | H | 30.68 | -94.09 | M | 19/06/2012 | SAMN55000511 |
| BALB05 | C | 10.05 | -83.33 | M | 31/03/2017 | SAMN55000481 |
| TN0421 | F | 36.55 | -87.14 | M | 24/08/2019 | SAMN55000525 |
| TN0102 | H | 35.03 | -89.35 | M | 22/08/2019 | SAMN55000524 |
| MO0312 | H | 36.56 | -90.35 | M | 18/08/2019 | SAMN55000506 |
| MI0125 | F | 42.23 | -84.49 | F | 28/08/2019 | SAMN55000471 |
| Il0411 | F | 42.02 | -89.33 | M | 14/08/2019 | SAMN55000496 |
| TT01.001 | H | 30.49 | -94.84 | M | 18/06/2012 | SAMN55000526 |
| LSSE05 | C | 10.42 | -84 | M | 27/03/2017 | SAMN55000499 |
| 11CT05 | D | 18.37 | -95 | M | 05/07/2011 | SAMN55000477 |
| 10OT10 | D | 18.68 | -96.38 | F | 08/10/2010 | SAMN55000459 |
| BZ5204 | D | 16.15 | -89.01 | M | 27/06/2021 | SAMN55000483 |
| NB0103 | A | 9.58 | -84.57 | M | 14/05/2014 | SAMN55000508 |
| 05AR27 | B | 18.96 | -103.95 | F | 26/07/2005 | SAMN55000461 |
| TM01_titiaB | G | 21.94 | -99.4 | M | 18/07/2004 | SAMN55000522 |
| IA0415 | F | 41.3 | -94.07 | M | 09/08/2019 | SAMN55000494 |
| LCBC04 | H | 31.17 | -89.58 | F | 26/08/2021 | SAMN55000470 |
| EAFB014150 | NA | 30.54 | -86.86 | M | 24/08/2021 | SAMN55000487 |
| CVa06 | H | 29.33 | -98.87 | M | 2021-06 | SAMN55000485 |
| ACSFc04 | E | 29.83 | -82.68 | M | 10/07/2019 | SAMN55000478 |
| SCSCa04 | E | 28.72 | -81.31 | M | 11/07/2019 | SAMN55000517 |
| NPKBa04 | H | 31.52 | -93.14 | F | 10/07/2017 | SAMN55000473 |
| RCJRa01 | F | 37.52 | -77.47 | M | 12/07/2019 | SAMN55000515 |
| SJ01.21 | H | 30.21 | -95.4 | M | 18/06/2012 | SAMN55000518 |
| OK0101 | H | 35.24 | -98.55 | M | 31/08/2019 | SAMN55000512 |
| MI0238 | F | 42.5 | -83.74 | M | 29/08/2019 | SAMN55000502 |
| CYa04 | G | 21.75 | -98.96 | M | 20/07/2004 | SAMN55000486 |
| TXRSa04 | D | 16.65 | -93.16 | M | 17/06/2021 | SAMN55000528 |
| MS0114 | H | 32.52 | -90.74 | M | 21/08/2019 | SAMN55000507 |

**Table S2.** Summary of general additive models (GAMs) of proportion of male wings with pigment, derived from manual measurements of hindwings from standard photos, as a function of measurements made on perched photos.

| **Proportion of hindwing pigmented (standard manual) ~ prop. of hindwing pigmented (manual measurements of perched individuals) (*n* = 66, Adj. R^2^= 0.98)** | | | | |
| --- | --- | --- | --- | --- |
| Parametric coefficients |  |  |  |  |
|  | Estimate | SE | *t* | *p*-value |
| (Intercept) | 0.676136 | 0.005126 | 131.9 | <0.001 |
| Smooth terms |  |  |  |  |
|  | edf | Ref.df | F | *p*-value |
| s(prop. hindwing pigmented [manual measurements of perched individuals]) | 2.912 | 2.994 | 936.7 | <0.001 |
| **Proportion of hindwing pigmented (standard manual) ~ prop. of hindwing pigmented (automated measurements of perched individuals) (*n* = 115, Adj. R^2^= 0.89)** | | | | |
| Parametric coefficients |  |  |  |  |
|  | Estimate | SE | *t* | *p*-value |
| (Intercept) | 0.4954 | 0.0106 | 46.73 | <0.001 |
| Smooth terms |  |  |  |  |
|  | edf | Ref.df | F | *p*-value |
| s(prop. hindwing pigmented [automated measurements of perched individuals]) | 2.939 | 2.997 | 322.8 | <0.001 |

**Table S3.** Summary of zero-one inflated beta model of proportion of male wings with pigment (*n* = 5892) as a function of an interaction between the day of year and population cluster (tree tip) (Bayesian R^2^ = 0.717, 95% CI = 0.710-0.724).

| **Parametric coefficients** |  |  |  |  |
| --- | --- | --- | --- | --- |
|  | Estimate | Est.Error | l-95% CI | u-95% CI |
| Intercept | -0.69 | 0.11 | -0.91 | -0.47 |
| phi_Intercept | 2 | 0.04 | 1.93 | 2.07 |
| zoi_Intercept | -6.72 | 0.69 | -8.27 | -5.58 |
| CLUSTERB | 0.26 | 0.16 | -0.05 | 0.59 |
| CLUSTERC | 1.35 | 0.18 | 0.98 | 1.69 |
| CLUSTERD | 1.73 | 0.15 | 1.44 | 2.03 |
| CLUSTERE | 1.83 | 0.16 | 1.52 | 2.14 |
| CLUSTERF | -0.95 | 0.17 | -1.32 | -0.65 |
| CLUSTERG | 1.33 | 0.15 | 1.04 | 1.62 |
| CLUSTERH | 1.03 | 0.12 | 0.79 | 1.27 |
| **Smooth terms** |  |  |  |  |
|  | Estimate | Est.Error | l-95% CI | u-95% CI |
| sds(sJulian.dateCLUSTERA_1) | 0.18 | 0.13 | 0.05 | 0.52 |
| sds(sJulian.dateCLUSTERB_1) | 0.35 | 0.17 | 0.15 | 0.81 |
| sds(sJulian.dateCLUSTERC_1) | 0.42 | 0.2 | 0.18 | 0.96 |
| sds(sJulian.dateCLUSTERD_1) | 0.74 | 0.31 | 0.37 | 1.54 |
| sds(sJulian.dateCLUSTERE_1) | 0.48 | 0.2 | 0.24 | 0.99 |
| sds(sJulian.dateCLUSTERF_1) | 0.07 | 0.09 | 0 | 0.31 |
| sds(sJulian.dateCLUSTERG_1) | 0.53 | 0.24 | 0.25 | 1.14 |
| sds(sJulian.dateCLUSTERH_1) | 0.58 | 0.23 | 0.31 | 1.15 |
| sds(phi_sJulian.dateCLUSTERA_1) | 2.36 | 0.85 | 1.2 | 4.46 |
| sds(phi_sJulian.dateCLUSTERB_1) | 5.53 | 1.55 | 3.35 | 9.26 |
| sds(phi_sJulian.dateCLUSTERC_1) | 0.4 | 0.29 | 0.02 | 1.12 |
| sds(phi_sJulian.dateCLUSTERD_1) | 1.86 | 0.65 | 0.97 | 3.5 |
| sds(phi_sJulian.dateCLUSTERE_1) | 1.32 | 0.49 | 0.7 | 2.55 |
| sds(phi_sJulian.dateCLUSTERF_1) | 1.43 | 0.86 | 0.53 | 3.58 |
| sds(phi_sJulian.dateCLUSTERG_1) | 0.38 | 0.37 | 0.01 | 1.36 |
| sds(phi_sJulian.dateCLUSTERH_1) | 1.01 | 0.39 | 0.52 | 1.98 |
| sds(zoi_sJulian.dateCLUSTERA_1) | 0.79 | 0.78 | 0.03 | 2.86 |
| sds(zoi_sJulian.dateCLUSTERB_1) | 0.65 | 0.63 | 0.02 | 2.41 |
| sds(zoi_sJulian.dateCLUSTERC_1) | 1.08 | 0.94 | 0.04 | 3.49 |
| sds(zoi_sJulian.dateCLUSTERD_1) | 6.26 | 1.99 | 3.53 | 11.21 |
| sds(zoi_sJulian.dateCLUSTERE_1) | 4.39 | 1.77 | 2.08 | 8.87 |
| sds(zoi_sJulian.dateCLUSTERF_1) | 1.88 | 1.97 | 0.05 | 6.65 |
| sds(zoi_sJulian.dateCLUSTERG_1) | 2.25 | 1.06 | 0.94 | 4.95 |
| sds(zoi_sJulian.dateCLUSTERH_1) | 7.14 | 3.74 | 2.81 | 16.95 |
| **Random effects** |  |  |  |  |
|  | SD est. | Est. error | l-95% CI | u-95% CI |
| Site.ID (intercept) | 0.52 | 0.03 | 0.46 | 0.57 |

**Table S4.** Leave-one-out cross-validation supports separate smooth terms for each cluster for the relationship between the day of year and μ, ϕ, and zero-one-inflation (zoi) terms in brms models. LOOIC is calculated as -2 * the expected log pointwise predictive density. Pareto-k diagnostics indicated acceptable reliability of the Pareto smoothed importance sampling, with only 0.7-1.3% of observations exceeding k = 0.7.

| **Model** | LOOIC | SE | ΔLOOIC |
| --- | --- | --- | --- |
| Full model (Supplementary Table 3)  μ ~  population cluster +  s(day of year* population cluster) +  (1 \| site.id)  ϕ ~  s(day of year * population cluster)  zoi ~  s(day of year * population cluster) | -6375.5 | 209.6 | 0 |
| Global smooth terms  μ ~  population cluster +  s(day of year) +  (1 \| site.id)  ϕ ~  s(day of year)  zoi ~  s(day of year) | -1674.9 | 174.3 | 4700.6 |
| Cluster specific smooth terms for mu only  μ ~  population cluster +  s(day of year * population cluster) +  (1 \| site.id)  ϕ ~  s(day of year)  zoi ~  s(day of year) | -2855.5 | 194.2 | 3520 |
| Cluster specific smooth terms for mu & phi only  μ ~  population cluster +  s(day of year * population cluster) +  (1 \| site.id)  ϕ ~  s(day of year * population cluster)  zoi ~  s(day of year) | -5256.6 | 219.5 | 1118.9 |
| Cluster specific smooth terms for mu & zoi only  μ ~  population cluster +  s(day of year * population cluster) +  (1 \| site.id)  ϕ ~  s(day of year)  zoi ~  s(day of year *population cluster) | -3978.0 | 182.9 | 2397.5 |

**Table S5.** Summary of zero-one inflated beta model of proportion of male wings with pigment, restricted to two-week windows with at least 10 observations (*n* = 5650), as a function of an interaction between the day of year and population cluster (tree tip) (Bayesian R^2^ = 0.713, 95% CI = 0.705-0.721).

| **Parametric coefficients** |  |  |  |  |
| --- | --- | --- | --- | --- |
|  | Estimate | Est.Error | l-95% CI | u-95% CI |
| Intercept | -0.68 | 0.18 | -1.03 | -0.32 |
| phi_Intercept | 1.84 | 0.05 | 1.74 | 1.94 |
| zoi_Intercept | -5.58 | 0.62 | -6.85 | -4.38 |
| CLUSTERB | 0.29 | 0.42 | -0.78 | 0.92 |
| CLUSTERC | 1.3 | 0.37 | 0.52 | 2 |
| CLUSTERD | 1.79 | 0.4 | 0.85 | 2.44 |
| CLUSTERE | 1.87 | 0.22 | 1.46 | 2.29 |
| CLUSTERF | -1.11 | 0.3 | -1.77 | -0.64 |
| CLUSTERG | 1.3 | 0.21 | 0.89 | 1.71 |
| CLUSTERH | 1.04 | 0.19 | 0.67 | 1.4 |
| **Smooth terms** |  |  |  |  |
|  | Estimate | Est.Error | l-95% CI | u-95% CI |
| sds(sJulian.dateCLUSTERA_1) | 0.27 | 0.26 | 0.04 | 0.96 |
| sds(sJulian.dateCLUSTERB_1) | 0.78 | 0.63 | 0.22 | 2.65 |
| sds(sJulian.dateCLUSTERC_1) | 0.63 | 0.39 | 0.23 | 1.69 |
| sds(sJulian.dateCLUSTERD_1) | 1.45 | 0.82 | 0.54 | 3.65 |
| sds(sJulian.dateCLUSTERE_1) | 0.59 | 0.3 | 0.26 | 1.3 |
| sds(sJulian.dateCLUSTERF_1) | 0.12 | 0.17 | 0 | 0.57 |
| sds(sJulian.dateCLUSTERG_1) | 0.55 | 0.26 | 0.24 | 1.25 |
| sds(sJulian.dateCLUSTERH_1) | 0.61 | 0.24 | 0.31 | 1.22 |
| sds(phi_sJulian.dateCLUSTERA_1) | 2.02 | 0.86 | 0.98 | 4.22 |
| sds(phi_sJulian.dateCLUSTERB_1) | 4.44 | 1.76 | 2.39 | 8.76 |
| sds(phi_sJulian.dateCLUSTERC_1) | 2.78 | 1.42 | 0.86 | 6.28 |
| sds(phi_sJulian.dateCLUSTERD_1) | 1.93 | 0.81 | 0.95 | 4.03 |
| sds(phi_sJulian.dateCLUSTERE_1) | 1.66 | 0.76 | 0.63 | 3.52 |
| sds(phi_sJulian.dateCLUSTERF_1) | 14.46 | 5.06 | 7.09 | 26.41 |
| sds(phi_sJulian.dateCLUSTERG_1) | 0.34 | 0.35 | 0.01 | 1.27 |
| sds(phi_sJulian.dateCLUSTERH_1) | 0.88 | 0.34 | 0.46 | 1.75 |
| sds(zoi_sJulian.dateCLUSTERA_1) | 2.3 | 2.52 | 0.07 | 8.94 |
| sds(zoi_sJulian.dateCLUSTERB_1) | 2.55 | 2.9 | 0.06 | 10.3 |
| sds(zoi_sJulian.dateCLUSTERC_1) | 1.3 | 1.29 | 0.06 | 4.47 |
| sds(zoi_sJulian.dateCLUSTERD_1) | 6.17 | 2.02 | 3.41 | 11.13 |
| sds(zoi_sJulian.dateCLUSTERE_1) | 3.73 | 1.63 | 1.59 | 7.83 |
| sds(zoi_sJulian.dateCLUSTERF_1) | 2.75 | 2.91 | 0.11 | 10.28 |
| sds(zoi_sJulian.dateCLUSTERG_1) | 1.88 | 0.96 | 0.73 | 4.4 |
| sds(zoi_sJulian.dateCLUSTERH_1) | 3.97 | 1.48 | 1.91 | 7.66 |
| **Random effects** |  |  |  |  |
|  | SD est. | Est. error | l-95% CI | u-95% CI |
| Site.ID (intercept) | 0.47 | 0.03 | 0.42 | 0.53 |

**Table S6.** Model fitting supports an OU, rather than a BM, model of trait evolution for both peak and off-peak wing phenotypes, with a higher stationary variance in the peak-season phenotypes.

| **Season** | **Model** | **phylogenetic variance (sigma)** | **alpha** | **stationary variance** | **AIC** |
| --- | --- | --- | --- | --- | --- |
| off-peak  (*n* = 1167) | BM | 0.40 | -- | -- | 2345.454 |
|  | OU | 0.12 | 14.28 | 0.004 | 2340.554 |
| peak (two-week window; *n* = 1015) | BM | 19.92 | -- | -- | 3427.818 |
|  | OU | 3.58 | 14.28 | 0.125 | 3420.802 |
| peak (one-week window; *n* = 427) | BM | 16.05 | -- | -- | 1497.313 |
|  | OU | 2.46 | 15.14 | 0.081 | 1489.176 |

**Table S7.** Locations and dates of phenotypic sampling for analyses of conspicuousness of melanized wings

| **Location Name** | **Drainage** | **Sampling Dates** | **Latitude** | **Longitude** |
| --- | --- | --- | --- | --- |
| Quebrada 20km | Pacific | 27 June – 05 July 2022 | 8.6186 | -83.0695 |
| Rio Sabalo | Pacific | 01 July – 06 July 2022 | 8.8681 | -83.3459 |
| Rio Carbón | Atlantic | 10 July 2022 | 9.6815 | -82.8343 |
| Cuba Creek | Atlantic | 11 July 2022 | 10.0117 | -83.2343 |
| San Miguel | Atlantic | 11 July 2022 | 10.0359 | -83.3270 |
| Rio Duruy | Atlantic | 12 July 2022 | 9.7072 | -82.9686 |

**Table S8.** Multi-state Markov models estimating transition rates (*q*) [mean + 95% CI] from ‘undamaged’ to ‘attacked’ as a function of region and season (*n* = 1515).

| **Model** | ***q*** | **AIC** | **ΔAIC** | **AIC.wi** |
| --- | --- | --- | --- | --- |
| Null (no covariates) | q_baseline = 0.0015  (0.0013, 0.0017) | 2925.462 | 67.117 | 0.000 |
| Regions | q_baseline = 0.0012  (0.0010, 0.0015)  q_northern= 0.2808 (0.1135,0.6947)  q_pacific = 2.545 (1.9,3.41) | 2861.854 | 3.509 | 0.105 |
| Season | q_baseline = 0.0015 (0.0013, 0.0017)  q_peak.season = 0.6265 (0.4724,0.8309) | 2916.845 | 8.500 | 0.000 |
| Region + Season | q_baseline = 0.0012  (0.0010, 0.0015)  q_northern= 0.3297 (0.1312, 0.8285)  q_pacific = 2.568 (1.917, 3.44)  q_peak.season = 0.747 (0.5605,0.9955) | 2859.851 | 1.506 | 0.286 |
| Region * Season | q_baseline = 0.0012  (0.0010, 0.0014)  q_atlantic_peak.season = 0.5198 (0.3189, 0.847)  q_northern_peak.season = 0.2153 (0.08581, 0.5404)  q_pacific_off.peak.season = 2.033  (1.392, 2.97)  q_pacific_peak.season= 1.877  (1.283, 2.745) | 2858.345 | 0.000 | 0.608 |

**Table S9.** Multi-state Markov models estimating transition rates (*q*) [mean + 95% CI] from ‘undamaged’ to ‘attacked’ as a function of region and season, excluding the Northern region (*n* = 1335).

| **Model** | ***q*** | **AIC** | **ΔAIC** | **AIC.wi** |
| --- | --- | --- | --- | --- |
| Null (no covariates) | q_baseline = 0.0017  (0.0015, 0.0019) | 2807.354 | 42.220 | 0.000 |
| Regions | q_baseline = 0.0015  (0.0013, 0.0018)  q_pacific = 2.545  (1.9,3.41) | 2768.643 | 3.509 | 0.105 |
| Season | q_baseline = 0.0017  (0.0015, 0.0019)  q_peak.season = 0.7684 (0.5766,1.024) | 2806.089 | 40.955 | 0.000 |
| Region + Season | q_baseline = 0.0015  (0.0013, 0.0018)  q_pacific = 2.568 (1.917,3.44)  q_season.peak = 0.747 (0.5605,0.9955) | 2766.64 | 1.506 | 0.286 |
| Region * Season | q_baseline = 0.0014  (0.0013, 0.0017)  q_atlantic_peak.season = 0.5197  (0.3189,0.8468)  q_pacific_off.peak.season = 2.033  (1.392, 2.969)  q_pacific_peak.season= 1.876 (1.283, 2.745) | 2765.134 | 0.000 | 0.608 |

**Table S10.** Multi-state Markov models estimating transition rates (*q*) [mean + 95% CI] from ‘undamaged’ to ‘attacked’ as a function of region and time to peak abundance of insectivorous birds, for deployments during the spring season (*n* = 675).

| **Model** | ***q*** | **AIC** | **ΔAIC** | **AIC.wi** |
| --- | --- | --- | --- | --- |
| Null (no covariates) | q_baseline = 0.0019 (0.0016, 0.0023) | 1570.207 | 27.038 | 0 |
| Region | q_baseline = 0.0018  (0.0015, 0.0022)  q_region_pacific = 2.033 (1.39, 2.97) | 1558.719 | 15.55 | 0 |
| Time to peak abundance | q_baseline = 0.0017  (0.0014, 0.0021)  q_time_to_peak = 0.7532 (0.6553,0.8658) | 1554.229 | 11.06 | 0.003 |
| Region + Time to peak abundance | q_baseline = 0.0016  (0.0013, 0.0020)  q_region_pacific = 2.011 (1.376,2.938)  q_time_to_peak = 0.75 (0.64,0.86) | 1543.169 | 0 | 0.725 |
| Region * Time to peak abundance | q_baseline = 0.0016 (0.0013, 0.0020)  q_region_pacific = 1.93 (1.07,3.49)  q_time_to_peak = 0.74 (0.60,0.90) q_region_pacific*time_to_peak= 1.03  (0.77,1.38) | 1545.136 | 1.967 | 0.271 |

**Table S11.** Multi-state Markov models estimating transition rates (*q*) [mean + 95% CI] from ‘undamaged’ to ‘attacked’ as a function of region and fold increase over the trough abundance during deployment (i.e., abundance at deployment/abundance at trough), for deployments during the spring season (*n* = 675).

| **Model** | ***q*** | **AIC** | **ΔAIC** | **AIC.wi** |
| --- | --- | --- | --- | --- |
| Null (no covariates) | q_baseline = 0.0019 (0.0016, 0.0023) | 1570.207 | 22.042 | 0 |
| Region | q_baseline = 0.0018  (0.0015, 0.0022)  q_region_pacific = 2.033 (1.39, 2.97) | 1558.719 | 10.554 | 0.004 |
| Fold increase in trough abundance at deployment | q_baseline = 0.0019  (0.0015, 0.0023)  q_fold_increase = 1.08 (1.01,1.16) | 1568.007 | 19.842 | 0 |
| Region + Fold increase in trough abundance at deployment | q_baseline = 0.0017  (0.0014, 0.0021)  q_region_pacific = 2.60 (1.69, 3.99)  q_fold_increase = 1.16  (1.06, 1.26) | 1549.857 | 1.692 | 0.299 |
| Region * Fold increase in trough abundance at deployment | q_baseline = 0.0017  (0.0013, 0.0021)  q_region_pacific = 13.22 (2.47, 70.75)  q_fold_increase = 1.18  (1.08, 1.29)  q_region_pacific*fold_increase= 0.682  (0.464,1.004) | 1548.165 | 0 | 0.697 |

**Table S12.** Multi-state Markov models estimating transition rates (*q*) [mean + 95% CI] from ‘undamaged’ to ‘attacked’ as a function of region and time to peak abundance of insectivorous birds, for deployments during the spring season and as a function of region and fold increase over the trough abundance during deployment (i.e., abundance at deployment/abundance at trough), for deployments during the spring season (*n* = 675). Analyses were conducted separately for attacks that were clearly made by birds (*n* = 118) and attacks that were clearly or likely made by birds (*n* = 141). In these analyses, attacks made by non-avian sources were treated as censored data points.

| **Model** | ***q*** | **AIC** | **ΔAIC** | **AIC.wi** |
| --- | --- | --- | --- | --- |
| **Clear bird attacks only: Time to peak abundance** | | | | |
| Null (no covariates) | q_baseline = 0.0010 (0.0008, 0.0013) | 885.3732 | 22.423 | 0.000 |
| Region | q_baseline = 0.0009 (0.0007, 0.0012)  q_region_pacific = 2.47 (1.45, 4.21) | 876.0659 | 13.1157 | 0.001 |
| Time to peak abundance | q_baseline = 0.0009 (0.0006, 0.0012)  q_time_to_peak = 0.69 (0.56, 0.85) | 872.287 | 9.3368 | 0.007 |
| Region + Time to peak abundance | q_baseline = 0.0008 (0.0006, 0.0011)  q_region_pacific = 2.48 (1.45, 4.22)  q_time_to_peak = 0.67 (0.54, 0.83) | 862.9502 | 0 | 0.701 |
| Region * Time to peak abundance | q_baseline = 0.0008 ( 0.0005, 0.0011)  q_region_pacific = 2.13 (0.95, 4.76)  q_time_to_peak = 0.63 (0.46, 0.88)  q_region_pacific*time_to_peak = 1.12 (0.72, 1.73) | 864.7094 | 1.7592 | 0.291 |
| **Clear bird attacks only: Fold increase in trough abundance at deployment** | | | | |
| Null (no covariates) | q_baseline = 0.0010 (0.0008, 0.0013) | 885.3732 | 19.0696 | 0.000 |
| Region | q_baseline = 0.0009 (0.0007, 0.0012)  q_region_pacific = 2.47 (1.45, 4.21) | 876.0659 | 9.7623 | 0.006 |
| Fold increase in trough abundance at deployment | q_baseline = 0.00010 (0.0008, 0.0013)  q_fold_increase = 1.09 (0.99, 1.20) | 884.5293 | 18.2257 | 0.000 |
| Region + Fold increase in trough abundance at deployment | q_baseline = 0.0008 (0.0006, 0.0012)  q_region_pacific = 3.47 (1.85, 6.51)  q_fold_increase = 1.21 (1.06, 1.37) | 869.6892 | 3.3856 | 0.154 |
| Region * Fold increase in trough abundance at deployment | q_baseline = 0.0008 (0.0006, 0.0011)  q_region_pacific = 47.99 (5.10, 451.90)  q_fold_increase = 1.25 (1.10, 1.42)  q_region_pacific*fold_increase = 0.54 (0.32,0.90) | 866.3036 | 0 | 0.839 |
| **Clear + likely bird attacks: Time to peak abundance** | | | | |
| Null (no covariates) | q_baseline = 0.0014 (0.0011, 0.0017) | 1142.965 | 12.988 | 0.001 |
| Region | q_baseline = 0.0013 (0.0010, 0.0017)  q_region_pacific = 1.68 (1.07, 2.65) | 1139.94 | 9.963 | 0.004 |
| Time to peak abundance | q_baseline = 0.0012 (0.0010, 0.0016)  q_time_to_peak = 0.76 (0.64, 0.89) | 1132.803 | 2.826 | 0.150 |
| Region + Time to peak abundance | q_baseline = 0.0012 (0.0009, 0.0015)  q_region_pacific = 1.67 (1.06, 2.62)  q_time_to_peak = 0.75 (0.63, 0.89) | 1129.977 | 0 | 0.617 |
| Region * Time to peak abundance | q_baseline = 0.0012 (0.0009, 0.0016)  q_region_pacific = 1.70 (0.83, 3.47)  q_time_to_peak = 0.75 (0.61, 0.94)  q_region_pacific*time_to_peak= 0.99 (0.70, 1.40) | 1131.973 | 1.996 | 0.227 |
| **Clear + likely bird attacks only: Fold increase in trough abundance at deployment** | | | | |
| Null (no covariates) | q_baseline = 0.0014 (0.0011, 0.0017) | 1142.965 | 13.07 | 0.001 |
| Region | q_baseline = 0.0013 (0.0010, 0.0017)  q_region_pacific = 1.68 (1.07, 2.65) | 1139.94 | 10.045 | 0.005 |
| Fold increase in trough abundance at deployment | q_baseline = 0.0013 (0.0010, 0.0016)  q_fold_increase = 1.11 (1.02, 1.20) | 1139.507 | 9.612 | 0.006 |
| Region + Fold increase in trough abundance at deployment | q_baseline = 0.0012 (0.0009, 0.0016)  q_region_pacific = 2.22 (1.33, 3.70)  q_fold_increase = 1.17 (1.06, 1.30) | 1131.969 | 2.074 | 0.259 |
| Region * Fold increase in trough abundance at deployment | q_baseline = 0.0012 (0.0009, 0.0015)  q_region_pacific = 18.34 (2.35, 142.90)  q_fold_increase = 1.20 (1.08, 1.32)  q_region_pacific*fold_increase= 0.61 (0.37, 0.98) | 1129.895 | 0 | 0.730 |

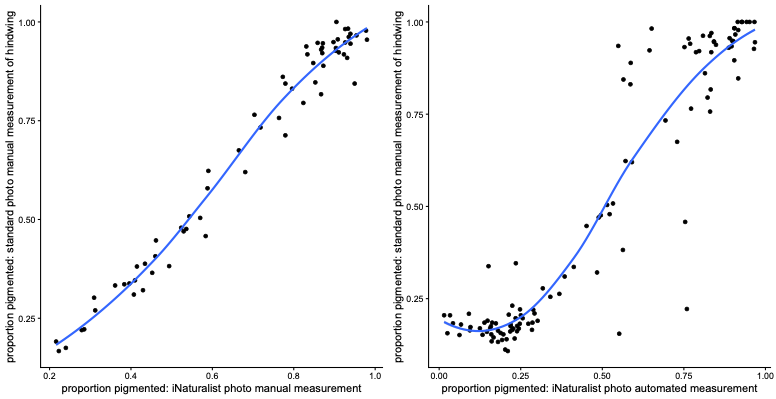

Fig. S1. Plots depicting the relationship between data derived from manual (left) and automated (right) measurements of iNaturalist photos and data derived from manual measurements of standard photos.

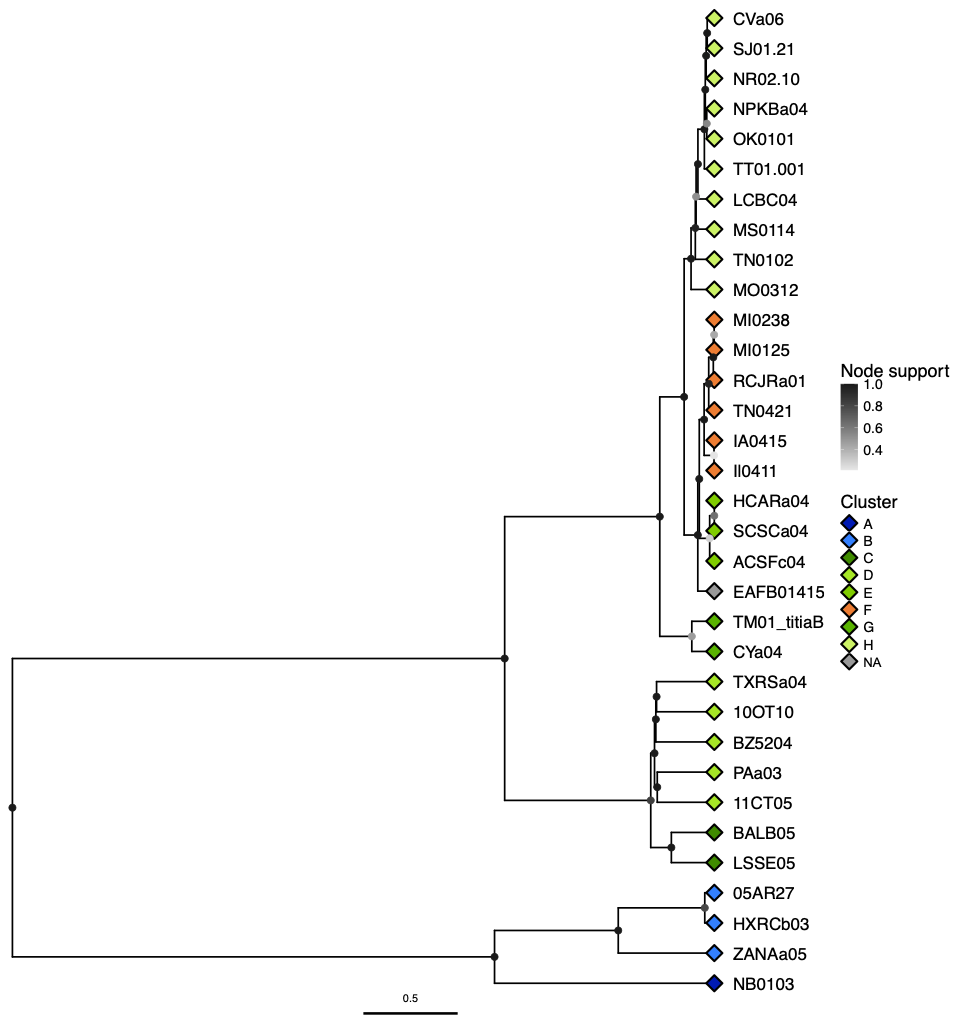
 **Fig. S2. Maximum clade credibility tree of whole-genome sequences of *Heaterina titia*.** Tip labels correspond to sample names in Table S1.

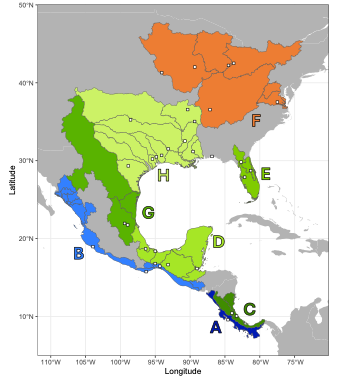

**Fig. S3. Plot of hydrobasins corresponding to each tip of the population-level phylogeny with points representing the sample locations of sequenced specimens.**

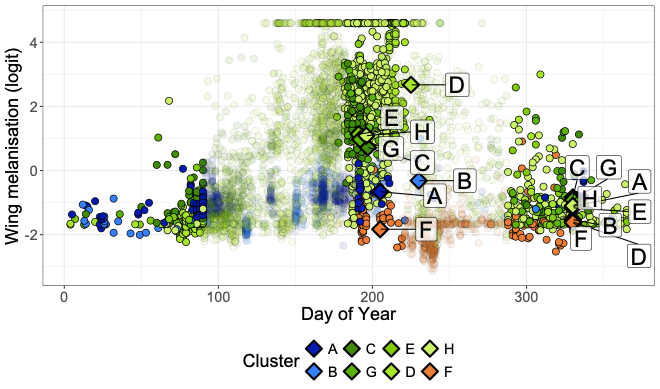

**Fig. S4. Logit-transformed values of wing pigmentation, with mean values for peak and trough measurements depicted for each tip of the population-level *Heaterina titia* phylogeny.**

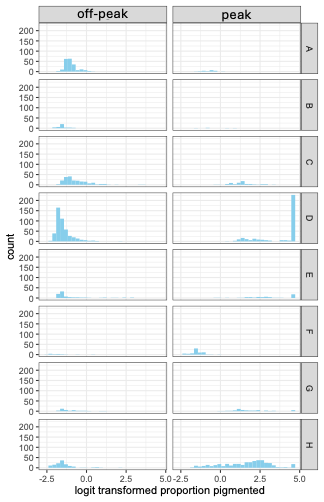

**Fig. S5. Histograms depicting the logit-transformed hindwing proportion pigmented for peak and off-peak measurements from each population cluster.**

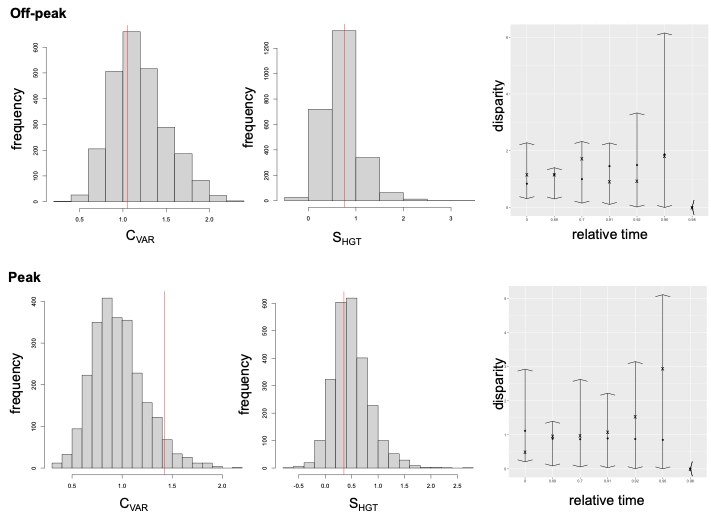

**Fig. S6. Model adequacy results for OU models fit to off-peak (top) and peak (bottom) measurements of wing pigmentation.** For C_VAR_ (left) and S_HGT_ (middle), the values for simulated datasets are depicted as histograms, with the values calculated on the empirical data depicted in red. For the rank envelope tests of disparity through time, the 95% CI for simulated datasets is depicted as lines, and the empirical values are depicted as Xs. In all cases, the empirical values were similar to simulated values (all *p* > 0.05).

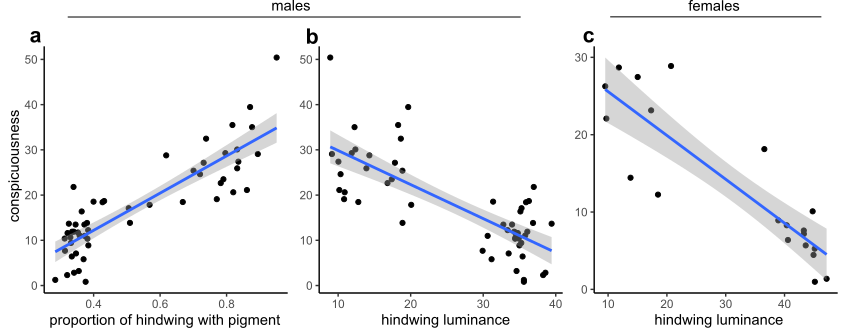

**Fig. S7. Increasing wing pigmentation is correlated with increasing conspicuousness.** Relationship between the **a.** extent and **b.** luminance of wing pigmentation and conspicuousness in males, and **c.** the luminance of wing pigmentation and conspicuousness in females.

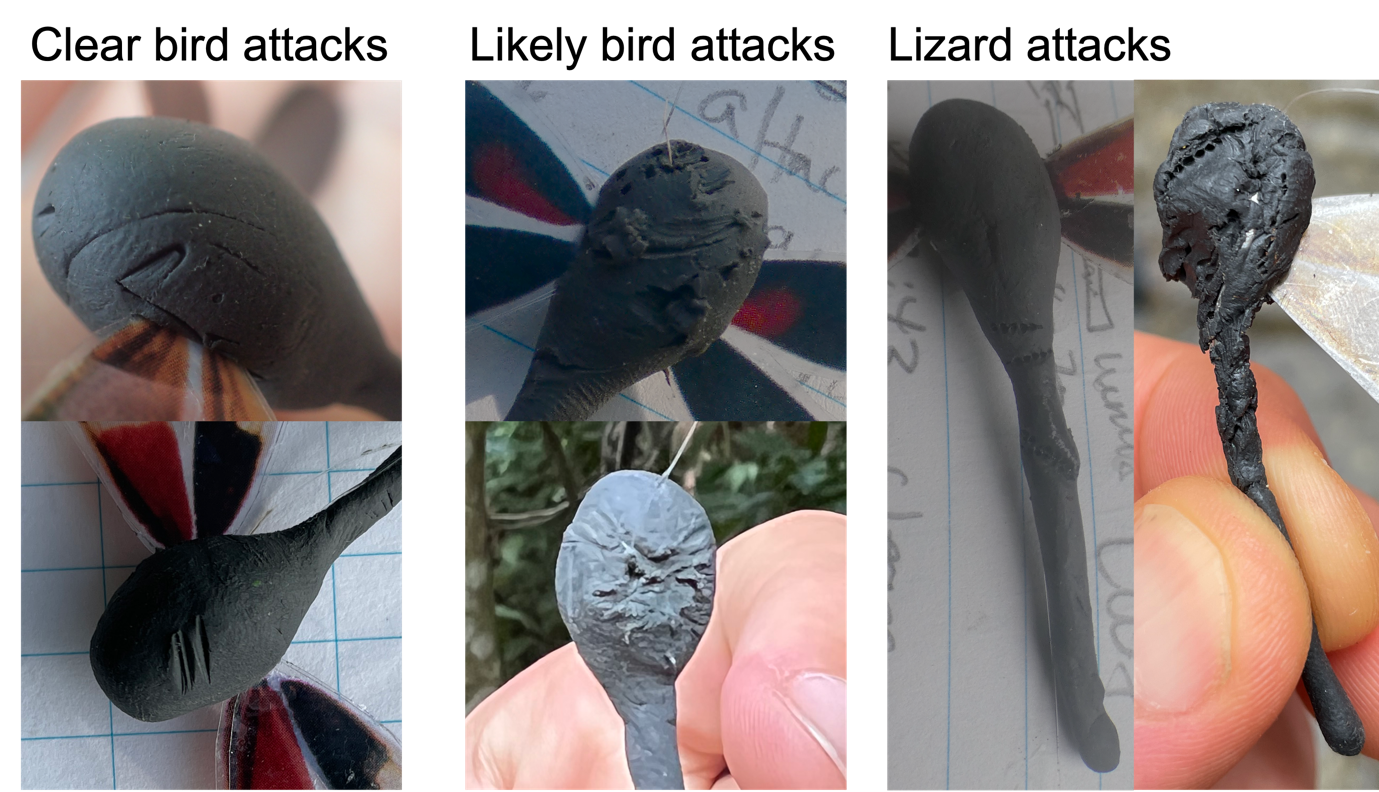

**Fig. S8. Examples of attacks on damselfly replicas.**
